## Supplementary Methods for "Broad diversity of human gut bacteria with traceable strain identifiers to facilitate bulk deposition"

Culture media

All quantities are given per liter of medium.

General process for agar plates: The components stated in the list below were supplemented with 15.0 g agar and a redox potential indicator (1.0 mg resazurin or 2.5 mg phenolsafranine). Afterwards they were dissolved in distilled water (99% of the final volume) and autoclaved (121°C, 15 min). A 100x concentrated stock solution containing heat sensitive components together with the reducing agents L-cysteine (final concentration 0.05% w/v) and DTT (final concentration 0.02% w/v) was created in distilled water and filter sterilized. All following steps were conducted in a laminar flow cabinet. After the autoclaved part of the media cooled down to approximately 60°C, the filtered stock solution was added (1% of the final volume). The still liquid agar was poured directly into petri dishes or one-well plates (Nunc™ OmniTray, Cat. No.: 140156). For the 96-well plates used in the case of single-cell dispensins, a multichannel pipette was used to transfer 250 µl of the medium into each well. The solidified medium was stored at 4°C and oxygen was removed by exposing the plates to an anaerobic environment for 48 h before use.

General process for Hungate tubes: The components stated in the list below were supplemented with a redox potential indicator (1.0 mg resazurin or 2.5 mg phenolsafranine) and the reducing agents L-cysteine (500 mg) and DTT (200 mg). After dissolving all components in distilled water to the final volume the medium was heated in a microwave until boiling and aliquoted into clean Hungate tubes (9 ml each). Oxygen was removed by submerging a gassing needle to the bottom of the Hungate and perfusing the hot medium with a forming gas/CO_2_ mixture (89.3% N_2_, 6% CO_2_, 4.7% H­_2_) for 3 min. While removing the needle the empty part of the Hungate tube was flushed with the gas mixture as well and immediately sealed with a rubber stopper and a screw cap. Finally the Hungate tubes were autoclaved (121°C, 15 min) and stored at room temperature until use.

List of media:

BBE (Bacteroides Bile Esculin with amikacin): (BD, ref. PA-254480.02) pancreatic digest of casein, 14.5 g; papaic digest of soybean meal, 5 g; NaCL, 5 g; esculin, 1 g; ferric ammonium citrate, 0.5 g; oxgall, 15 g; hemin, 0.01 g; amikacin 0.075 g; vitamin K1, 0.01 g; growth factors 1.8 g.

BHI (Brain-heart-infusion): (DSM Medium 215c) BHI (Oxoid, ref. CM1135B), 37 g.

GMM (Gut Microbiota Medium): tryptone peptone, 2 g; yeast extract, 1 g; D-glucose, 0.4 g; L-cystein, 0.5 g; cellobiose, 1 g; maltose, 1 g; fructose, 1 g; meat extract, 5 g; KH_2_PO_4_, 13.6 g; MgSO_4_ x 7 H_2_O, 0.002 g; NaHCO_3_, 0.4 g; NaCl, 0.08 g; CaCl_2_, 0.008 g; vitamin K (menadione), 0.001 g; FeSO_2_, 0.4 mg; hematin, 1.2 mg solved in 0.2M histidine; Tween80, 0.05 % v/v; ATCC Trace Mineral Mix, 10 ml; acetic acid, 1.7 ml; isovaleric acid, 0.1 ml; propionic acid, 2 ml; butyric acid, 2 ml. Heat sensitive components (added after autoclaving): ATCC Vitamin Mix, 10 ml.

mGAM (modified Gifu Anaerobic Medium): (DSM Medium 1715) GAM Agar, Modified (HyServe, ref. 05433), 41.7g.

mGAM blood: mGAM as described above with addition of defibrinated sheep blood, 50 ml. The sheep blood was added after autoclaving (without filter sterilization) and only used for the creation of agar plates.

WCA (Wilkins-Chalgren Anaerobe broth): (DSM Medium 339a) WCA (Oxoid, ref. CM0643), 33 g.

YCFA (Yeast Casitone Fatty Acids): (DSM Medium 1611) Casitone, 10.0 g; yeast extract, 2.5 g; glucose, 5.0 g; MgSO_4_ x 7 H_2_O, 0.045 g; CaCl_2_ x 2 H_2_O, 0.09 g; K_2_HPO_4_, 0.45 g; KH_2_PO_4_, 0.45 g; NaCl, 0.9 g; NaHCO_3_, 4.0 g; hemin, 0.01 g; acetic acid, 1.9 ml; propionic acid, 0.7 ml; isobutyric acid, 90.0 µl; n-valeric acid, 100.0 µl; iso-valeric acid, 100.0 µl. Heat sensitive components (added after autoclaving): Biotin, 0.02 mg; folic acid, 0.02 mg; pyridoxine-HCl, 0.1 mg; thiamine-HCl x 2 H_2_O, 0.05 mg; riboflavin, 0.05 mg; nicotinic acid, 0.05 mg; D-Ca-pantothenate, 0.05 mg; vitamin B_12_, 10.0 µg; p-aminobenzoic acid, 0.05 mg; lipoic acid, 0.05 mg.
