## Supplementary Results for "Broad diversity of human gut bacteria with traceable strain identifiers to facilitate bulk deposition"

Variability in genome size and functionality between phyla

Genome size variation was observed between phyla, with *Actinomycetota* genomes being the smallest (2.38 ± 0.35 Mbp). *Bacteroidota* had the largest genomes (4.97 ± 1.02 Mbp), although not significantly larger than *Pseudomonadota* (4.84 ± 0.64 Mbp; adj. *p* = 0.56) (**Supplementary Figure** **4a**; **Supplementary Table** **3**). This variation in genome size between phyla was validated using >5,000 previously published genomes ^1^ (**Supplementary Figure 4b/c; Supplementary Table** **3**). In addition to having the largest genomes, *Bacteroidota* also had the largest repertoire of CAZymes (**Supplementary Figure 5b**). To account for the variation in coding potential of each strain, and thereby examine their investment in carbohydrate utilisation, we normalised their CAZyme load based on their number of coding sequences (**Supplementary Figure 5c**). This analysis confirmed that the *Bacteroidota* have the highest investment in carbohydrate utilisation. However, they were followed by the *Actinomycetota* (**Supplementary Table** **4**), indicating that, whilst members of *Actinomycetota* have undergone host-associated genome size reduction ^2^, they have invested in carbohydrate degradation.

16S rRNA gene sequence-based ecological analysis

To assess the occurrence of HiBC strains in the human gut, we looked at their prevalence and relative abundance within 1,000 human gut samples (**Supplementary Figure** **6**). This analysis was complemented by screening the genome of each isolate against a collection of 42,927 MAGs ^1,3^. *Bifidobacterium* spp. were some of the most prevalent (~70% of samples) and abundant (~9% on average) species. Cultivation and sequencing approaches capture different fractions of the human gut; this is highlighted by our ability to isolate a strain of *Selenomonas* *noxia*, which was only observed in a single sample, with a relative abundance <0.1%, making it a sub-dominant member of the gut microbiota, likely of oral origin ^4^. Although only present at an average relative abundance of 2.7%, strains of *Agathobacter* *rectalis* (synonym *Eubacterium rectale*) were identified as being the most frequently reconstructed strain (n = 1,839 MAGs). In contrast, 19 strains did not match any MAGs, including seven novel taxa. To further assess the importance of HiBC species to the human gut ecosystem, we screened them against the original Human Microbiome Projects “most wanted” list of taxa ^5^. Of the 1,468 most wanted sequences, 90 matched HiBC strains, including 4 high, 24 medium, and 62 low priority sequences. Of these, 8 matching strains represented novel taxa within the phylum *Bacillota* according to the current state of nomenclature, including two novel genera (**Supplementary Figure 7**). These “most wanted” species, and all other novel taxa discovered in this study are described in the main text at the end of the methods.

Effect of plasmids on ANI value calculation

Given the number of plasmids a single strain can contain, and the size a single plasmid can reach (see main results), we tested whether they could affect the taxonomic identification of an isolate, as most genomes lack delineation of plasmids. Given the promiscuous nature of plasmids ^6^, it is possible that the taxonomic signal of a strain may be masked by the presence of multiple large plasmids, distorting taxonomic assignment. As average nucleotide identity (ANI) is a standard approach for taxonomic assignment of genomes, we calculated the theoretical plasmid size required to potentially cause a >1% change in ANI between two genomes of identical size (**Supplementary Figure 8a**). Plasmids >80 kp should theoretically cause such a difference in ANI within genomes ≤8 Mb. To assess this within the HiBC strains, we selected all plasmid-containing strains that had a close relative (determined by GTDB-Tk) (n = 152). We then calculated the ANI to these close relatives, both with and without plasmid inclusion (**Supplementary Figure 8b**). The plasmids had a significant (*p* = 0.01), but negligible (delta-ANI, 0.004 ± 0.01) effect on reducing the ANI to the closest relative strains. The effect of plasmid inclusion on ANI was independent of cumulative plasmid size, with some small plasmid sets having a greater effect on ANI than the largest (**Figure 2d**). The only strain on which the removal of plasmids had a significant effect was CLA-ER-H4, which was assigned to 'Collinsella sp900547855' by GTDB-Tk when its plasmids were included, but was assigned to 'Collinsella sp003487125' without the plasmids. This is indicative of the need for better taxonomic study of this genus, as the plasmids had no effect on the assignment of CLA-ER-H4 to 'Collinsella sp003487125', but removing the plasmids increased ANI to 'Collinsella sp900547855' by 0.05%. However, to avoid the creation of false species within *Collinsella*, we have not described any new species within this genus, which needs to be done in a comprehensive manner.

Ecology of the pMMCAT megaplasmid

The matching pMMCAT plasmids were observed within species of *Bacteroides*, *Phocaeicola* and *Parabacteroides* from the USA, China, Japan, and Israel (**Supplementary Figure 1c**). Network analysis of the pMMCAT plasmids suggested those from *Phocaeicola* were more divergent than the *Bacteroides* and *Parabacteroides* pMMCAT plasmids (**Supplementary Figure 1d**). MobMess^15^ analysis assigned pMMCAT_H253 as a compound plasmid, supporting the theory that it contains additional cargo, leading to the variable associations with health conditions for the predicted proteins. A sequence within the pMMCAT family was detected in 99.55% (1,774/1,782) of human gut metagenomes ^15^. However, only 52 metagenomes had detection values >95%, including two samples from Fiji, with the highest coverage (135.82x) being from USA samples (**Supplementary Figure 1e**).

Bulk submission system at the DSMZ

The DSMZ welcomes bulk deposits (>10 isolates) ([www.dsmz.de/bulk-deposit](http://www.dsmz.de/bulk-deposit)) of microbial strains isolated from host species or environmental habitats. Bulk deposits are time-intensive and take considerably longer to process than single strains. We therefore encourage depositors to start bulk deposits as early as possible in their projects to have them ready before the submission of accompanying manuscripts. Requests for urgent processing of bulk deposits during the revision phase of a manuscript cannot be considered.

Bulk deposits are processed in a standardised 3-step procedure.

1. Depositors are asked to fill out the [Excel Bulk Deposit Accession Form](https://www.dsmz.de/fileadmin/user_upload/Deposit/DSMZ_Bulk_Deposit_Form_2024_06_28.xlsx) covering all relevant strain information and sign the [Bulk Deposit Agreement](https://www.dsmz.de/fileadmin/user_upload/Deposit/Agreement_Bulk_Deposit_DSMZ.pdf).
2. The coordinating curator requests the shipment of a first batch of up to 10 test strains for quality and legal checks as required for deposition in the DSMZ collection.
3. If all strains of step 2 pass these checks, the coordinating curator requests the remaining strains to be subsequently shipped in batches of 30-40 strains at a time. After the processing of each batch is completed, the next batch will be requested until the entire bulk deposit has been processed.

Depositors are strongly encouraged to request for integration of their strains into StrainInfo, which provides DOIs early in the deposition process (e.g., to use them in publications and sequence submissions). This enhances the traceability of the strains throughout the deposition process. In this case the responsible curator can approve integration of a culture and its identity data into StrainInfo once the culture has been received by the DSMZ and passed initial checks.  At this point the strain is assigned a DOI and is published on StrainInfo with the status “Deposition in progress”. Once the deposition process of the strain is completed the public status of the strain in the database is changed to “Published” with online catalogue status “available”.
